# Augmented Interval List: a novel data structure for efficient genomic interval search

**DOI:** 10.1101/593657

**Authors:** Jianglin Feng, Aakrosh Ratan, Nathan C. Sheffield

**Affiliations:** UNIVERSITY OF VIRGINIA

## Abstract

**Motivation:** Genomic data is frequently stored as segments or intervals. Because this data type is so common, interval-based comparisons are fundamental to genomic analysis. As the volume of available genomic data grows, developing efficient and scalable methods for searching interval data is necessary.

**Results:** We present a new data structure, the augmented interval list (AIList), to enumerate intersections between a query interval *q* and an interval set *R*. An AIList is constructed by first sorting *R* as a list by the interval *start* coordinate, then decomposing it into a few approximately flattened components (sublists), and then augmenting each sublist with the running maximum interval *end*. The query time for AIList is *O*(*log*_2_*N* + *n* + *m*), where *n* is the number of overlaps between *R* and *q, N* is the number of intervals in the set *R*, and *m* is the average number of extra comparisons required to find the *n* overlaps. Tested on real genomic interval datasets, AIList code runs 5 - 18 times faster than standard high-performance code based on augmented interval-trees (AITree), nested containment lists (NCList), or R-trees (BEDTools). For large datasets, the memory-usage for AIList is 4% - 60% of other methods. The AIList data structure, therefore, provides a significantly improved fundamental operation for highly scalable genomic data analysis.

**Availability:** An implementation of the AIList data structure with both construction and search algorithms is available at code.databio.org/AIList.

## 1. Introduction

A genomic interval *r* is defined by the two coordinates that represent the *start* and *end* locations of a feature on a chromosome. The general interval search problem is defined as follows:

Given a set of *N* intervals *R* ={*r*_1_, *r*_2_, *…, r*_*N*_} for *N*≫1, and a query interval *q*, find the subset *S* of *R* that intersect *q*. If we define all intervals to be half-open, *S* can be represented as:

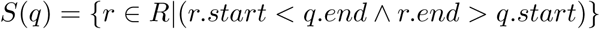

If the order of intervals in *R* remains the same when the elements are sorted based on interval *start* or based on interval *end*, then *S* can be computed using a single binary search followed by another *n* + 1 comparisons, where *n* is the number of overlaps between *R* and *q*. We refer to *R* in this case as being *flat*. The strategy of a binary search followed by sequential comparisons becomes sub-optimal when intervals in *R* possess a coverage or containment relationship, *i.e.*, one interval covers or contains another interval in *R*. Such non-*flat* interval lists require extra comparisons beyond *n* + 1 after the binary search.

The interval search problem is fundamental to genomic data analysis (Giardine, 2005; Li and Durbin, 2009; Layer *et al.*, 2018; Jalili *et al.*, 2018) and several approaches have been developed to do this efficiently (Cormen *et al.*, 2001; Kent *et al.*, 2002; Richardson, 2006; Alekseyenko and Lee, 2007; Quinlan and Hall, 2010; Neph *et al.*, 2012). Currently, the most popular data structures are the nested containment list (NCList) by Alekseyenko and Lee (2007), the augmented interval tree (AITree) by Cormen *et al.* (2001), and the R-tree by Kent *et al.* (2002). The design of these data structures can be conceptualized as minimizing the additional comparisons in the search strategy we described earlier. NCList and AITree require *O*(*Nlog*_2_*N*) time to build the data-structures, and their query time complexities are *O*(*log*_2_*N* + *n* + *m*), where *m* is the average number of extra comparisons (comparisons that do not yield overlapping results) required to find the *n* overlaps. The R-tree has a complexity of *O*(*N*) in construction and *O*(*m* + *n*) in query. For genomic interval datasets, *N* is usually several orders of magnitudes larger than *n* or *m*.

NCLists, AITrees and R-trees differ both in time to build the data structure and to search it. As shown in Alekseyenko and Lee (2007), both average construction time and search time can vary significantly among the methods in practice. The search time differences are determined by the extra comparison value *m*, which differs based on the different approaches of each algorithm: For NCList, many sublists may be involved in a query, which requires many extra binary searches; for AITree, one must compare all interval nodes marked by the augmenting value, and not all of them intersect the query; and for R-tree, all intervals in an indicated bin are scanned, although many of these may not overlap the query.

In this paper, we present a new data structure, the Augmented Interval List (AIList). The AIList search algorithm reduces the number of extra comparisons (*m*) and thereby achieves better performance.

## 2. AIList data structure and query algorithm

### 2.1 A simple augmented interval list and query algorithm

To begin, we sort *R* based on the interval *start* to create an interval list. We then augment the list with the running maximum *end* value, *MaxE*, which thus stores the maximum *end* value among all preceding intervals (Figure 1a) to create the AIList. *MaxE* reflects the containment relationship among the intervals: because the 2nd interval [3, 8) contains the 3rd interval [5, 7), they have the same *MaxE* value (8); similarly the 4th interval [7, 20) contains the next three intervals and they all have the the same *MaxE* value 20. *MaxE* therefore indicates a list of containment groups. The coverage length of a containment group can be defined as the number of sequential identical *MaxE* values minus one. A variation of this approach instead augments the list with the sorted end value, *SortedE*, which we detail in the Supplementary Information.

**Fig. 1:**
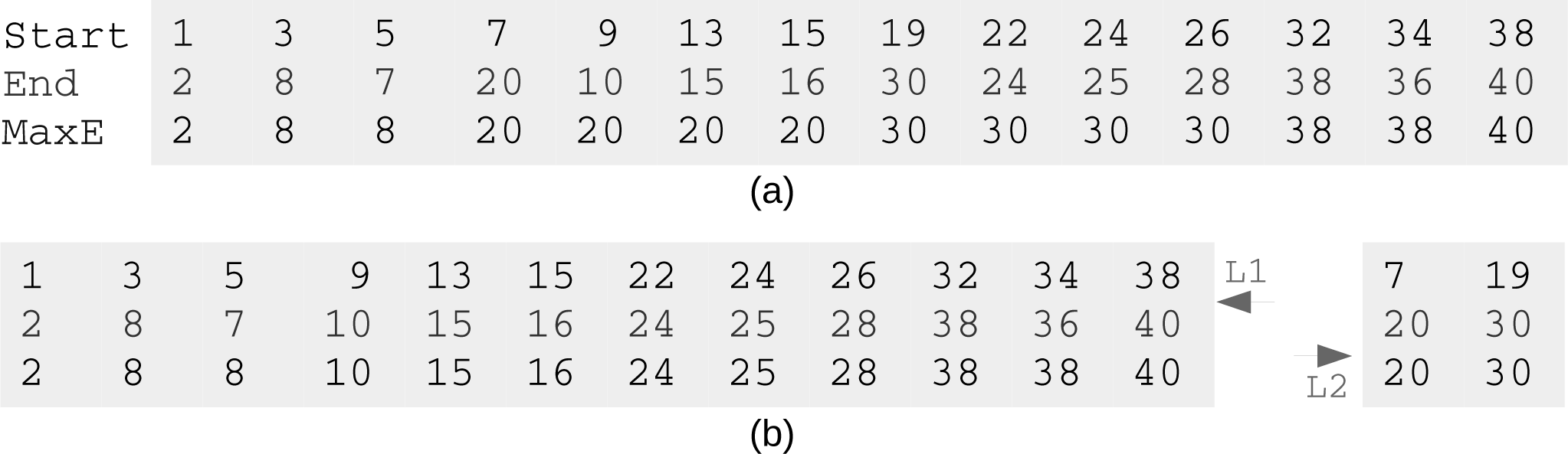
*(a) An interval list sorted by the `Start` and augmented with `MaxE`, the maximum `end` counting from the first interval. (b) Interval list decomposition: Intervals in the above list with `End` value larger than that of the three following intervals are put into a separated list. The two component lists L*1 *and L*2 *are both flattened. Two queries [`9,12`) and [`17,21`) are discussed in the text.*

Once an AIList is constructed, we seek to search it with a query interval. To find the overlaps of a query [*q.start, q.end*) with the AIList, we first do a binary search using the query interval *end* against the sorted list of database interval *starts* to find the right-most interval in the AIList where *r.start* < *q.end*. We then step backwards to test for each interval whether *MaxE > q.start*. Since *MaxE* is in ascending order, and *MaxE* ≥*r.end*, when we find the first interval with *MaxE* ≤*q.start*, we are guaranteed that all remaining intervals do not overlap the query and can be skipped. For example, if our query were [9, 12), we first binary search to find the index of the last interval *I*_*E*_ that has *r.start* < 12, which is the 5th interval [9, 10), *MaxE* = 20. We know all intervals on the right side will not overlap the query since their *r.start* ≥12. We then step backwards to the 5*th*, then the 4*th* interval, but we can stop at the 3*rd* interval, [5, 7) with *MaxE* = 8, since *MaxE* < 9, which ensures that no further intervals on the left side will overlap the query. This is very efficient since we only checked one extra interval.

However, this strategy becomes less efficient in the presence of long coverage groups. For example, if the query were [17, 21), then we need to check 6 intervals: from the 8th, [19, 30) with *MaxE* = 30, back to the 3rd, [5, 7) with *MaxE* = 8, which requires 4 extra comparisons. This example query is the worst case for this list. To mitigate the issue of containment, we next extend the AIList by decomposing the list.

### 2.2 Decomposition of an interval list

Because long coverage groups lead to extra comparisons, we seek to reduce the length of these groups. We achieve this by extracting containing intervals into a second list. In the simple case shown in Figure 1a, we can define the coverage length *len* of a list interval *i* as the number of the immediately following intervals (*i* + 1, *i* + 2, …) that are covered by interval *i*; so in Figure 1a *len*[1] = 0, *len*[2] = 1, *len*[4] = 3, etc. Then, we can extract all intervals that have coverage length larger than or equal to a criterion (minimum coverage length *MinL*). In the above example, we can set *MinL* = 3 to decompose the list into two sublists, *L*1 and *L*2 (Figure 1b). Then we add *MaxE* to each *L*1 and *L*2 independently and attach *L*2 to the end of *L*1 to form an improved augmented interval list (AIList) with the same size as the original list. The start of the sublists in the new AIList are maintained in a header list *hSub*. Now in *L*1 there are only two containment groups, both of length 1, and there is no containment in *L*2.

#### Algorithm 1 AIList Construction Algorithm

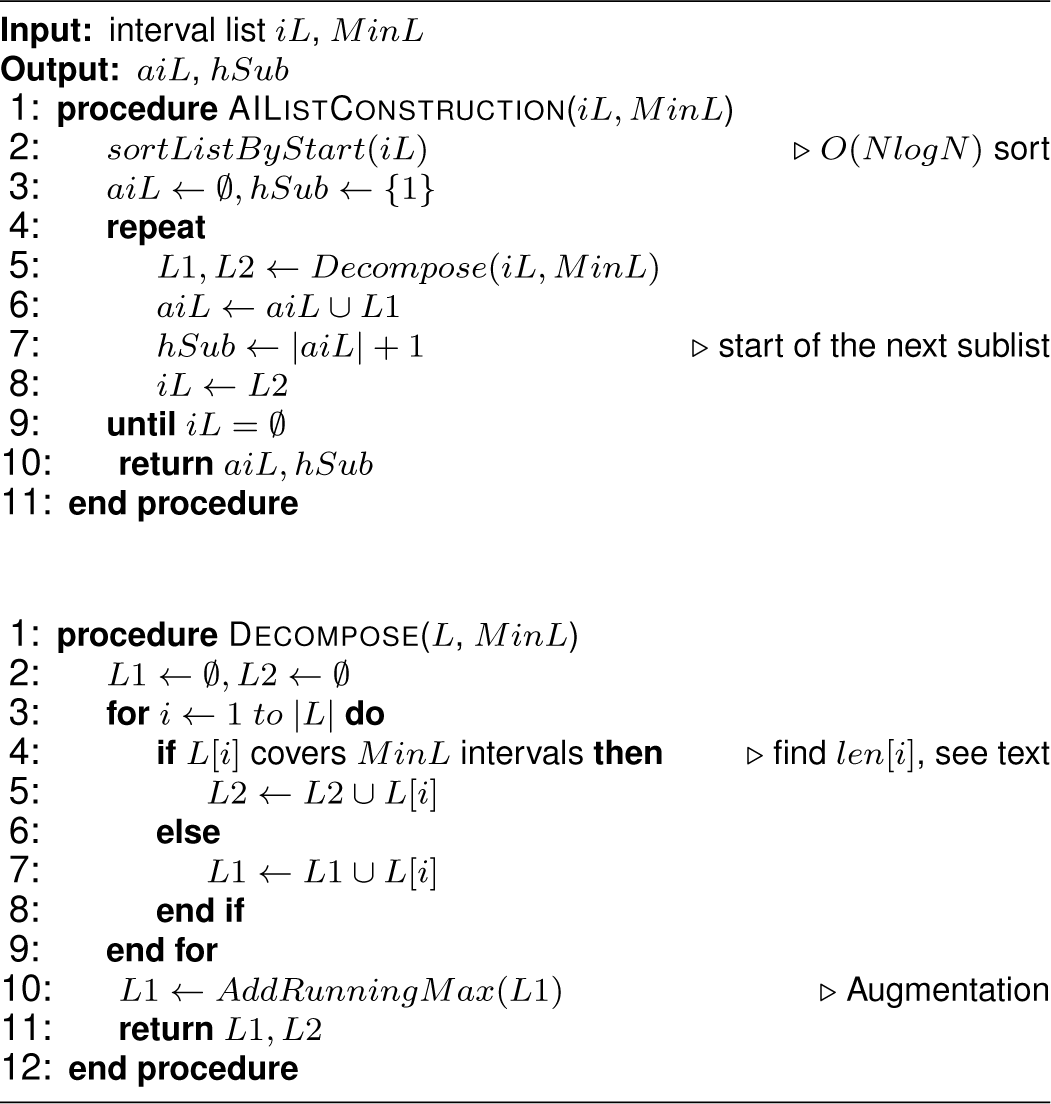

For a more complex dataset, the *L*2 sublist may in turn have long containments, in which case the decomposition is repeated recursively as long as *L*2 is decomposable. The practical implementation of this decomposition may need to relax our definition of the coverage length *len* to include more general cases. For example, if *iL*[5] does not contain *iL*[6], but it contain all intervals from *iL*[7] to *iL*[12], we may still want to extract *iL*[5]. The AIList data structure can accommodate different ways to implement the decomposition depending on how *len* is defined. In our implementation, we define coverage length *len*[*i*] for interval item *i* as the number of intervals among the next 2 ∗ *MinL* intervals that are covered by interval *i*. Therefore, to find *len*[*i*] we can simply check intervals from *iL*[*i* + 1] to *iL*[*i* + 2 ∗ *MinL*] to count how many of them are covered by *iL*[*i*]. Limiting the search range to 2 ∗ *MinL* reduces construction time.

Selection of *MinL* determines the extent of decomposition, with greater values leading to less decomposition. The optimal *MinL* differs mildly among datasets (Fig. 2). It is clear from Figure 2 that *MinL* = 20 is near optimal for datasets we tested, and more importantly, that the runtime is relatively robust to substantial variation in *MinL* choice, so it is not necessary to find the optimal *MinL* for each dataset. We have set the default *MinL* to 20, which results in the number of components *nSub* being less than 10 for all the datasets we have tested (see Table 1).

**Table 1.**
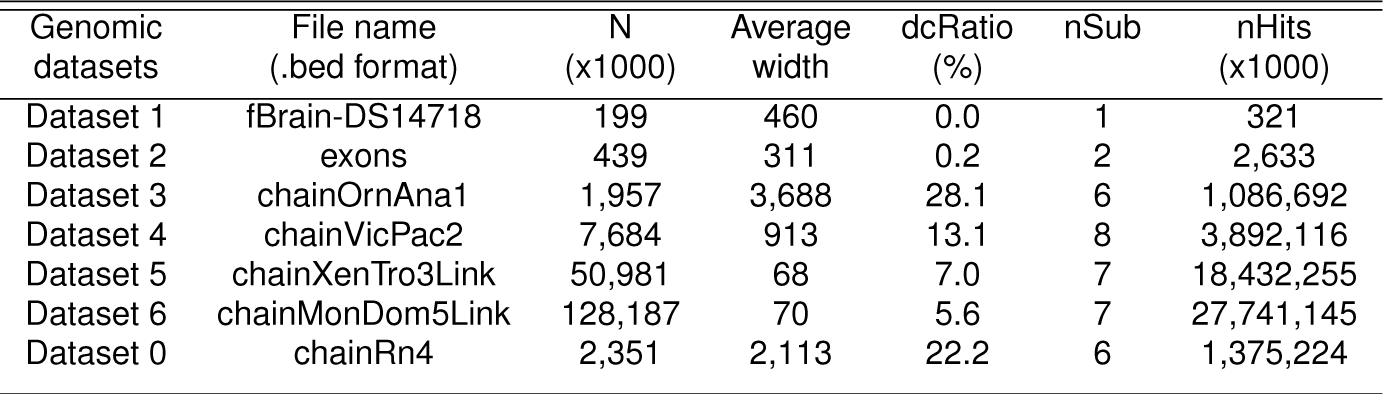
Genomic interval datasets used as database or query for performance evaluation. *N* is the number of intervals, *dcRatio* is the ratio of the number of intervals that cover their immediate next over *N*, *nSub* is the number of total sublists for AIList, *nHits* is the number of overlaps with query Dataset 0. Dataset 1 and 2 are from BEDTools, and others are from UCSC.

| Genomic datasets | File name<br>(.bed format) | N<br>(x1000) | Average<br>width | dcRatio<br>(%) | nSub | nHits<br>(x1000) |
| --- | --- | --- | --- | --- | --- | --- |
| Dataset 1 | fBrain-DS14718 | 199 | 460 | 0.0 | 1 | 321 |
| Dataset 2 | exons | 439 | 311 | 0.2 | 2 | 2,633 |
| Dataset 3 | chainOrnAna1 | 1,957 | 3,688 | 28.1 | 6 | 1,086,692 |
| Dataset 4 | chainVicPac2 | 7,684 | 913 | 13.1 | 8 | 3,892,116 |
| Dataset 5 | chainXenTro3Link | 50,981 | 68 | 7.0 | 7 | 18,432,255 |
| Dataset 6 | chainMonDom5Link | 128,187 | 70 | 5.6 | 7 | 27,741,145 |
| Dataset 0 | chainRn4 | 2,351 | 2,113 | 22.2 | 6 | 1,375,224 |

**Fig. 2:**
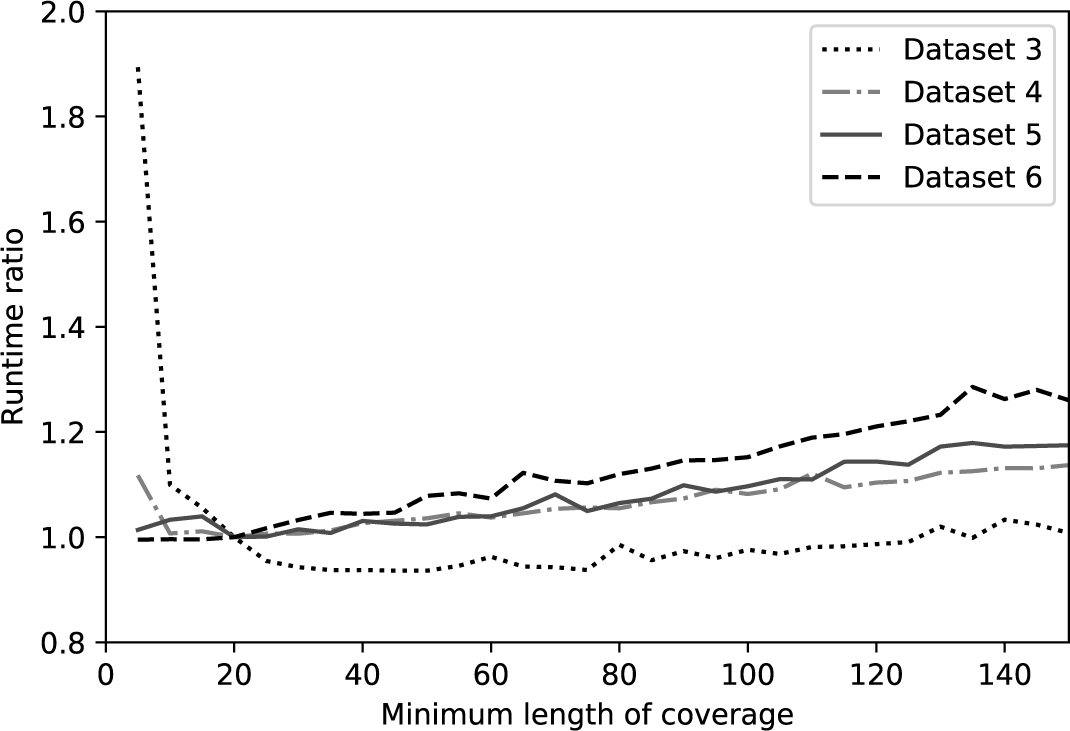
*Runtime ratio as a function of `MinL` value for four different datasets with a fixed query dataset. Since the tuntime for the four different sized datasets differ significantly, each curve is scaled to the runtime of its own at `MinL`* = 20.

Algorithm 1 lists a simplified *O*(*NlogN*) algorithm for constructing an AIList, including both decomposition and augmentation. The function *AddRunningMax* is a simple linear scan to determine the *maxE* value.

### 2.3 The AIList query algorithm including decomposed sublists

Queries against the decomposed list structure are similar to the original case, but now done independently on each sublist. The decomposition process has divided the original list into 2 or more flattened or nearly flattened sublists, so queries in each sublist are close to optimal. The cost of this improvement is that we now require additional binary searches to get the indices *I*_*E*_ for each sub-list. This approach thus implements a tradeoff between number of binary search comparisons and number of extra interval comparisons due to interval containment. Similar to other algorithms mentioned above, the query time complexity for AIList is *O*(*log*_2_*N* + *n* + *m*), but the average number of extra comparisons *m* is minimized. The search algorithm is listed in Algorithm 2.

#### Algorithm 2 AIList Search Algorithm

**Fig. 2:**
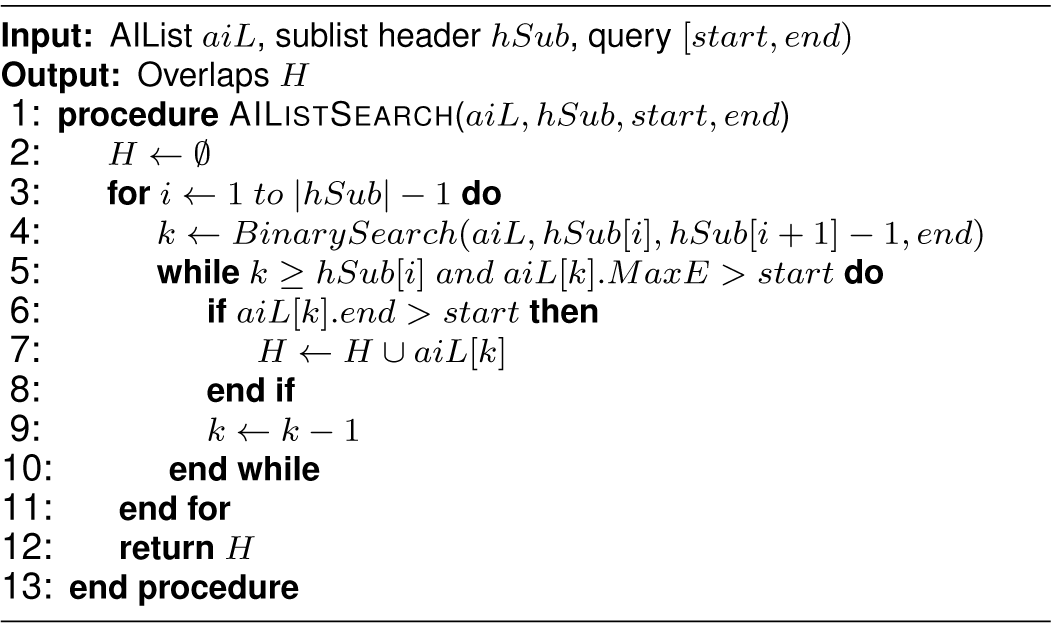

## 3. Results

AIList is implemented in C. To evaluate the efficiency of AIList, we compared its performance with interval trees (AITree), nested containment lists (NCList), and R-trees. For AITree we used the rbtree-based interval tree from Linux kernel (see Supplementary Information); for NCList we used the C-code *intervaldb.c* implemented by Alekseyenko and Lee (2007); and for R-tree we used the popular C++ implementation BEDTools v2.25.0 by Quinlan and Hall (2010). AIList outperforms other approaches on all datasets that we have tested so far. Table 1 lists seven real and representative genomic datasets ranging in size from 199,000 to 128 million intervals. In each of the experiments we describe below, we used Dataset 0 as the *query* set, with the other six used as the *database* interval set. All experiments were run on a computer with 2.8GHz CPU and 16GB memory, and the reported runtime for each method includes the time required for data loading, data structure construction, searching, and result output.

For all datasets, AIList outperformed all other methods (Table 2). The improvement was more dramatic for the datasets with more complex containment structure: For *flat* dataset 1 and near *flat* dataset 2,

**Table 2.**
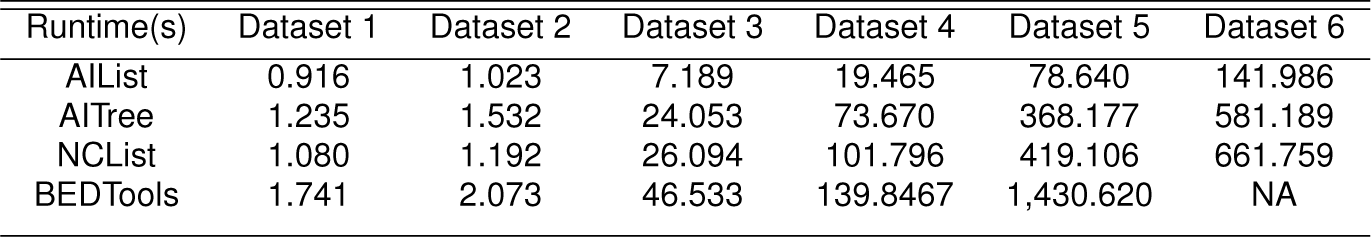
*Runtime (seconds) of AIList, AITree, NCList and BEDTools for datasets listed in Table 1. Dataset 1 to 6 are used as database and dataset 0 is as query set. No result for BEDTools on dataset 6 since it took nearly all of the machine memory (16GB) and was terminated.*

AIList is 120-150% faster than AITree and NCList and 2 times faster than BEDTools. For datasets with greater containment, AIList is up to 5 times faster than AITree and NCList, and up to 18 times faster than BEDTools. Furthermore, AIList consumed substantially less memory than the other algorithms (Table 3).

**Table 3.** *Memory usage (%, memory used by a program divided by the total machine memory) of AIList, AITree, NCList and BEDTools for large datasets on a computer with a total memory of 16 GB. BEDTools was terminated for Dataset 6 because it took all machine memory.*

| Dataset | AIList | AITree | NCList | BEDTools |
| --- | --- | --- | --- | --- |
| Dataset 5 | 3.7 | 19.6 | 6.2 | 95.4 |
| Dataset 6 | 9.2 | 49.2 | 19.1 | NA |

To evaluate how AIList, AITree, NCList and R-tree scale in practice for differing sizes of input dataset, we selected a single comparison (Dataset 0 queried against Dataset 5 as input dataset) and then downsampled the input dataset in increments of 1,000,000 intervals. Fig. 3a shows four curves of runtime vs input dataset size. Fig. 3b shows the ratio of the runtime compared to AIList. As input dataset size increases from 1 million to 10 million, the runtime ratio increases from 2.8 to 8.3 for AITree, from 2.8 to 4.5 for NCList, and from 6.9 to 20 for BEDTools, demonstrating the superior scaling of AIList.

**Fig. 3:**
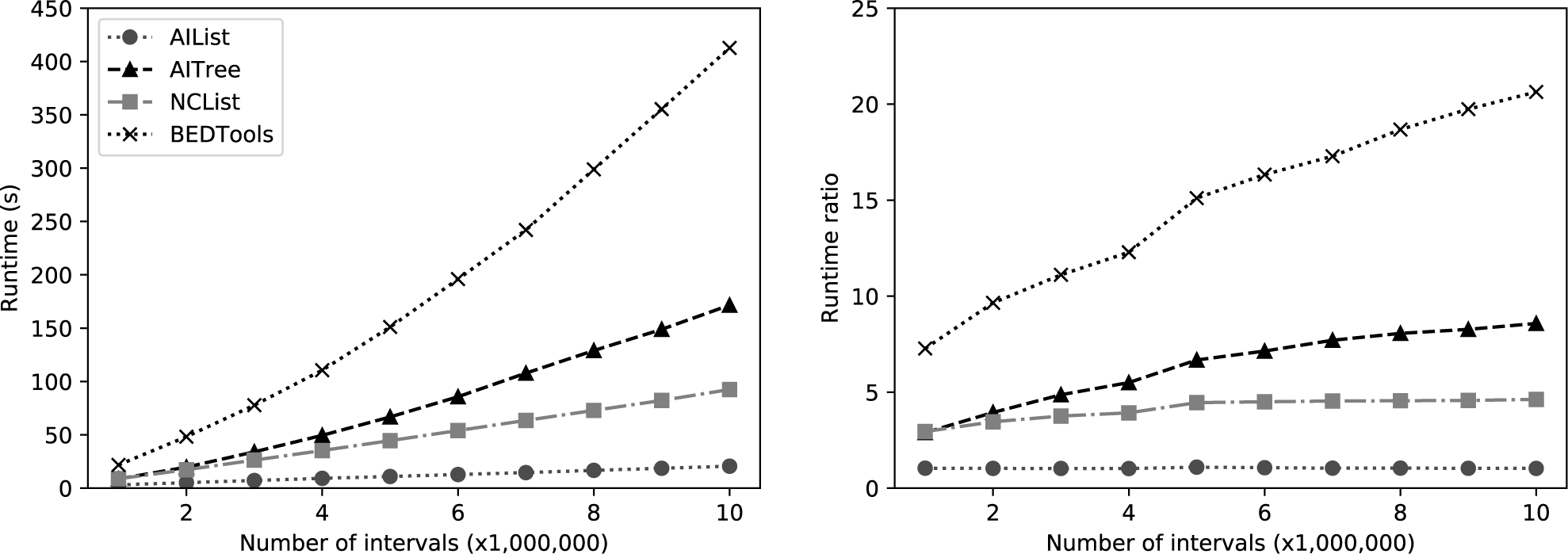
*Performance comparison of AIList with AITree, NCList and BEDTools as a function of the size of the target dataset. Dataset 0 is the query set, target datasets are subsets of Dataset 5 by sampling.*

We also performed a similar simulation experiment by subsampling the query, which demonstrates the practically lower scaling of the AIList algorithm (Figure 4). As expected, all algorithms scale linearly but AIList has the lowest coefficient.

**Fig. 4:**
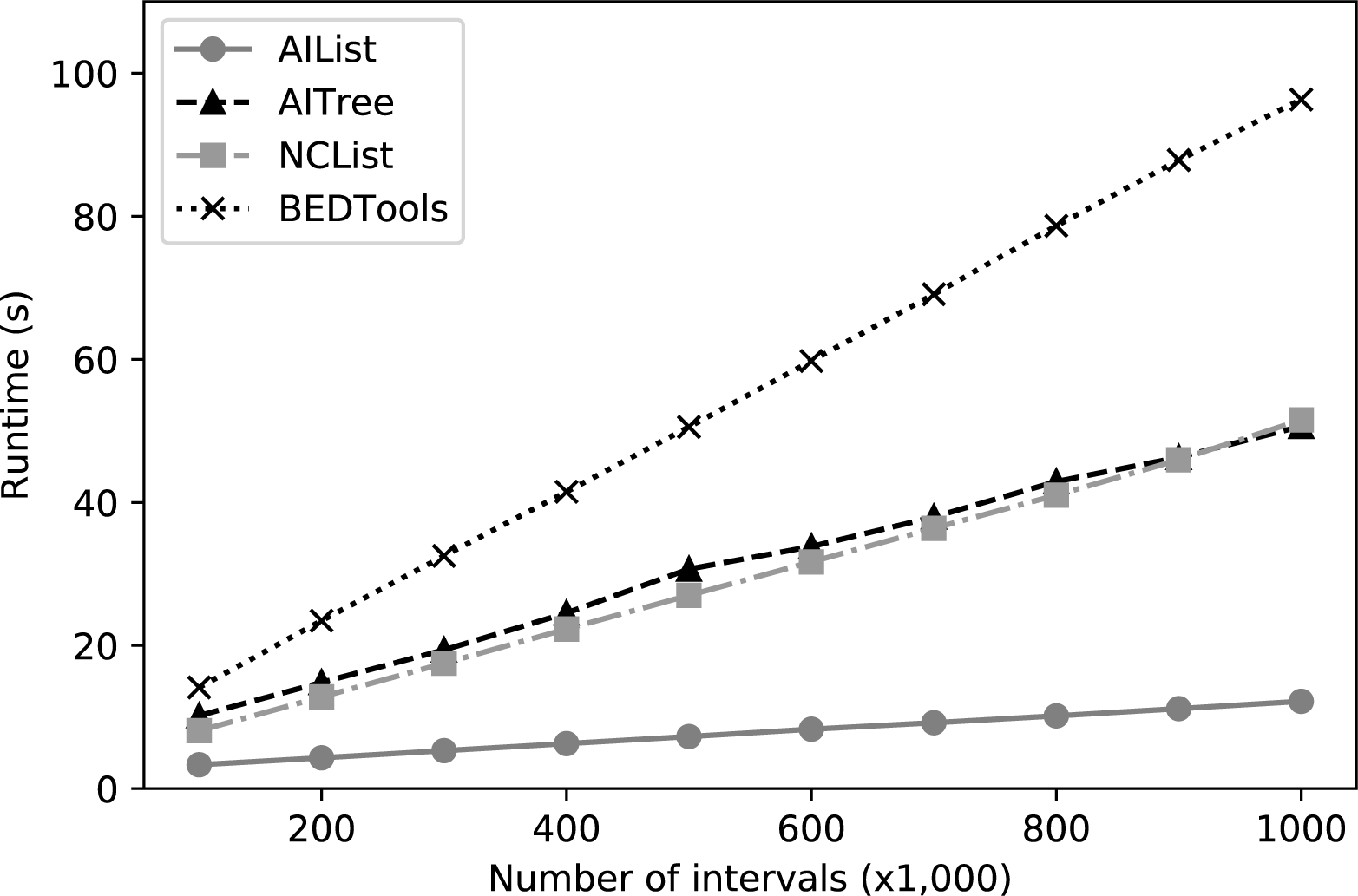
*Performance comparison of AIList with AITree, NCList and BEDTools as a function of the size of the query interval set. Dataset 4 is the target dataset, query sets are subsets of Dataset 0 by sampling.*

## 4. Conclusion

We defined a novel data structure that dramatically improves on the enumeration of interval overlaps. Our method pairs the techniques of list augmentation and list decomposition to provide tighter terminal conditions for overlap checking, which reduces the number of extra comparisons that must be made. We demonstrated that this method outperforms existing methods across a series of datasets that range from flat structure to highly nested interval containment.

Similar to the NCList, the AIList data structure contains only one extra data element *MaxE*, so it is more efficient than the AITree (3 extra elements) and the R-tree (contains duplicate elements); but the AIList header size is negligible, while the NCList header size can be comparable to database size (see Supplementary Information for details). Another advantage of the AIList is that inserting a new interval element requires simply checking the few sublists to find roughly where it belongs, which is much more flexible than NCList, which can require reconstructing the whole data structure. Because of its simple data structure, AIList is also the simplest to implement (see Algorithm 1 and 2, and source code). Finally, as shown in Table 3, AIList takes the least memory.

Taken all these together, AIList provides a significantly improved fundamental operation for highly scalable genomic data analysis.

